# “Multigenerational effects of temperature exposure on upper thermal limit and mitochondrial functioning in Medaka (*Oryzias Latipes*) brain”

**DOI:** 10.1101/2025.03.09.642098

**Authors:** Julie Morla, Mélissa Richard, Rémy Lassus, Nicolas Pichaud, Martin Daufresne, Arnaud Sentis

## Abstract

Determining the mechanisms underpinning the thermal limits of organisms is essential to anticipate the impact of climate change on their survival. As mitochondrial activity is key for cellular ATP production, we aimed at investigating the link between mitochondrial dysfunction and the upper thermal limit (CT_max_) at which neuromuscular functions are lost. We reared medaka fish (O*ryzias latipes)* over generations at 20°C (COLD) and 30°C (WARM) during five years and measured their CT_max_ and their mitochondrial oxygen consumption rates (OCRs) at 25°C and at their CT_max_. We found that multigenerational exposure at elevated temperature increases the CT_max_ by 13%. We observed for all fish a general decrease in the OCRs and for the RCR of complex-I between 25°C and their CT_max_ suggesting a link between CT_max_ and cerebral mitochondrial performance. Finally, we found an increase in the contribution of the mitochondria complex II to OCRs at elevated temperature suggesting that this complex underlie the plasticity in mitochondrial functioning at high temperature. Our study highlights the link between whole organism thermal tolerance and mitochondrial dysfunction and that multigenerational exposure can modify mitochondrial and thermal performance.

## Introduction

Temperature is a fundamental environmental factor highly affecting metabolic and physiological processes, through the increased speed of biochemical reactions in ectotherms. Global warming and the expected increase in the frequency of extreme thermal events (IPCC 2023) is thus imposing strong constraints on ectothermic organisms. A rapid increase in temperature, particularly in shallow aquatic environments that are highly sensitive to temperature fluctuations, is a determining physiological stress factor for aquatic ectotherms (Woodward, 2009). In the current context, a better understanding of the thermal tolerance of ectotherms is thus crucial to predict the vulnerability of aquatic ectotherms to these rapid changing environmental conditions.

Thermal vulnerability indices such as safety margins and warming tolerance have increased in popularity for predicting species vulnerability to climate change (Clusella-Trullas et al., 2021). For each species, we can define specific tolerated temperature ranges in which optimum temperature values represent maximum performance (T°Opt) as well as temperatures at which performance is negatively affected and where the species survival is jeopardized. Cowles and Bogert in 1944 defined the upper *limit* to thermal tolerance as “*the thermal point at which locomotor activity becomes disorganized and the animal loses its ability to escape from conditions that will promptly lead to its death*” (Cowles and Bogert, 1944). Such *limit is generally refers as to the* critical thermal maximal (CT_max_; Martínez et al., 2016; O’Donnell et al., 2020). CT_max_ is considered as an excellent proxy of upper thermal tolerance as it trigger similar behavioural responses across a diversity of species (Lutterschmidt and Hutchison, 1997).

Understanding the limiting physiological processes determining the CT_max_ has been the focus of several previous studies (reviewed in Chung and Schulte, 2020). For most organisms, thermal responses are determined by physiological processes at the metabolic level. As detailed by Chung and Schulte., (2020), in many species the fact that patterns of genetic variation in the mitochondrial genome are correlated with environmental temperature has led to the formulation of the “mitochondrial climate adaptation hypothesis” (Camus et al., 2017). This hypothesis suggests that mitochondria may impose limits on whole-organism thermal tolerance. Following the Arrhenius law, the metabolism of ectotherms increases with warming, accelerating cellular respiratory and oxygen demand (Arrhenius, 1889; Hilton et al., 2010; Sommer and Pörtner, 2004). In a recent review, Ern et al., (2023) reported that rapid direct thermal impacts on fish act through three fundamental molecular mechanisms: *reaction rates, protein structure, and membrane fluidity*. They stipulate that during acute warming, these molecular effects lead to loss of equilibrium/death through variety of pathways including mitochondrial dysfunction. They also summarized that the increased energy demands of ectotherms exposed to heat, combined with the molecular mechanisms affected, make the failure of mitochondrial processes a likely mechanism underlying whole organism thermal failure.

Mitochondria are central to cellular respiration as they use oxygen to transduce energetic substrates (nutrients) into adenosine triphosphate (ATP), needed to maintain essential body functions. Mitochondrial respiration is determined as the respiration supporting ATP synthesis by oxidative phosphorylation (OXPHOS, the coupling between substrate oxidation and ATP synthesis) and the respiration required to offset proton in the absence of ATP synthesis (LEAK). Enzyme and protein complexes of the respiratory chain that allow cellular respiration are profoundly influenced by temperature (see appendix Fig. S1; Abele et al., 1998; Clotfelter et al., 2013; Somero et al., 2017). Previous studies reported that the different mitochondrial complexes have different thermal sensitivities and may play a role in thermal adaptation of ectotherms (Blier et al., 2014; Chung and Schulte, 2020). In particular, the first complex (complex I) of the respiratory chain has been suggested to be a site of thermal mitochondrial injury after a critical heat exposure and could be the main determinant of heat tolerance (El-Wadawi and Bowler, 1996; Jørgensen et al., 2021; Menail et al., 2022). In addition, the failure of mitochondrial oxygen consumption for complex I is associated with an increase in the production of signalling molecules that can cause damage in excessive amounts (Reactive Oxygen Species, i.e. ROS) and enhance macromolecular damages (Genova et al., 2004; Jørgensen et al., 2021; Lenaz et al., 1997). Thus, the study of these complexes can help to explain the thermal limits of ectotherms, which holds important significance in the context of the current climate warming and species adaptation.

According to Chung and Schulte., (2020), data from many species suggest that there may be a link between mitochondrial failure and failure of higher-level processes such as brain failures, cardiac failure and whole-organism thermal limitations (Chung and Schulte, 2020; Ekström et al., 2016; Fangue et al., 2009; Harada et al., 2019; Hraoui et al., 2020; Martínez et al., 2016). Mitochondria may specifically mediate heart failure under heat stress and play a role in CT_max_ (Ekström et al., 2016; Hunter-Manseau et al., 2019; Iftikar et al., 2014; Iftikar and Hickey, 2013; Pichaud et al., 2017). However, mitochondrial dysfunction could also underlie neuronal failure. Indeed, mitochondrial oxygen consumption measured in the brain could indicate greater mitochondrial thermal sensitivity compared to cardiac tissue in *Fundulus heteroclitus* (Chung et al., 2017).

Early, neuronal system dysfunction induced by temperature have been suggested to constrain whole-organism thermal limits (Ern et al., 2015; Somero and DeVries, 1967). Friedlander et al., (1976), demonstrated a functional link between brain function and whole-animal thermal tolerance reporting that brain failure occurs at the same temperature as the whole organism in goldfish (*Carassius auratus*) (Friedlander et al., 1976). Comparable results are reported in Atlantic cod (*Gadus morhua*), and zebrafish (*Danio rerio*) (Andreassen et al., 2022; Jutfelt et al., 2019). Recent studies have thus tested the hypothesis that the loss of equilibrium during acute warming is due to direct thermal effects on neuronal function in the brain (Temperature-Dependent deterioration of Electrical Excitability, TDEE hypothesis) (Robertson, 2004; Vornanen, 2020). While this hypothesis may contribute to explain whole-organism thermal limits, recent studies concluded that it is probably not the only mechanisms leading to heat failure (Ern et al. 2023).

Indeed, the link between mitochondrial failure and failure of higher-level processes is not well established and remains a controversial topic issue (Bilyk and DeVries, 2011; Clark et al., 2013; Ern et al., 2015; Schulte, 2015; Ørsted et al., 2022). Early studies showed little evidence of a functional link between the mitochondrial function and the whole organism thermal tolerance because in vitro rates of mitochondrial oxygen consumption were maintained at temperatures above the CT_max_ (Somero et al., 1996; Somero, 2002; Pörtner, 2002; Chung and Schulte, 2020). Based on previous studies, Ern et al., (2023) proposed that more accurate assessments of mitochondrial function can be achieved using the respiratory control ratio (RCR) (i.e. an estimate of mitochondrial efficiency) and state III respiration (an estimate of the maximal capacity of mitochondria to produce ATP) (Brand and Nicholls, 2011; Chance and Williams, 1955). RCR is determined as the ratio of the oxygen consumption rate supporting ATP production to the oxygen consumption rate required to offset proton leakage (Brand and Nicholls, 2011; Chance and Williams, 1955). Based on these measures, Chung and Schulte., (2020) reported a correlation between acute thermal limits of ectotherms and the acute limits of mitochondrial function. It is however important to consider that these proxies of mitochondrial functions entail important bias, as they are often measured with excessive amounts of substrates and specific substrate types (e.g. NADH-linked such as pyruvate and malate, only fuelling complex I). Indeed, mitochondrial complexes can use other substrates and notably succinate (FADH2-linked to fuel complex II) which denotes of a certain flexibility in substrates oxidation. As a matter of fact, it has been shown that this flexibility is especially important at high temperatures, at which a switch from NADH-linked to FADH_2_-linked substrates are often observed albeit not in all ectotherms (Jorgensen et al., 2021; Menail et al., 2022; Blanchard et al., 2024; Hraoui et al., 2021; Hraoui et al., 2020; Chung et al., 2017; Willis et al., 2021; Hickey et al., 2024).

To provide more evidence of this relationship, Ern et al., (2023) suggested to preferentially use thermal acclimation to manipulate in concert the thermal tolerance of the mitochondrial components and the whole animal, a concept that has been recently applied in Drosophila (Blanchard et al., 2024). Several studies suggest that CT_max_ varies with the exposure duration and may thus be enhanced after a period of acclimation to elevated temperature (Angilletta, 2001; Bilyk and DeVries, 2011). Some studies found that thermal acclimation can lead to plastic modifications of the structure and function of mitochondria in ectotherms (Guderley and St-Pierre, 2002; Seebacher et al., 2010). Warm acclimation generally results in a decrease in mitochondrial metabolism at elevated temperatures, due to a decrease in mitochondrial respiration rates, or a decrease in mitochondrial density or through compositional changes within mitochondrial membranes (Ekström et al., 2016; Schulte, 2015; Yan and Xie, 2015; Chung et al., 2017; Morla et al., 2024; Blanchard et al., 2024). These adjustments may result both from reversible phenotypic plasticity, notably observable in mitochondria, and from genetic adaptation over generations (Guderley and St-Pierre, 2002; Seebacher et al., 2010) and can have an impact on population performance. In a recent study, Rank et al. 2020, reported that population of montane leaf beetle can exhibit latitudinal differences in mitochondrial and nuclear genotypes that are associated with differences in performance traits, suggesting that mitochondrial thermal limits can affect whole organism performance. Due to the complexity of the thermal acclimation process, the different components of mitochondrial metabolism affected by temperature are unclear although it can be crucial to predict and anticipate the impacts of rapid global warming on ectotherms. To go further, an effective strategy to identify the mechanisms underlying thermal tolerance is to study thermal responses after multiple generations of exposures. Multiple generations can produce lineages that differ in their innate thermal tolerances or in their thermal acclimation capacity (Morgan et al., 2020).

In this study, our objective was to determine multigenerational temperature exposure effects on (1) the upper critical thermal maximum (CT_max_) of the whole organism medaka fish (*Oryzias latipes*), (2) the oxygen consumption rates, OCRs in brain mitochondria and (3) a potential link between a dysfunction of brain mitochondria and whole organism CT_max_. We hypothesized that (1) brain mitochondria can limit the whole organism CT_max_ and (2) that CT_max_ can increase after a multigenerational exposure at elevated temperature resulting in mitochondrial changes (Angilletta, 2001; Bilyk and DeVries, 2011; Somero and DeVries, 1967; Ern et al., 2015). To test our hypotheses, we assessed the CT_max_ of medaka fish maintained over several generations to cold (20°C) and warm (30°C) temperatures by exposing fish of each thermal origin to a progressive increase in temperature until it loses its locomotor function. In a second step, we measured the mitochondrial OCRs of fish from the two lines at 25°C and at their mean CT_max_ focusing on differences between the contributions of the different mitochondrial complexes to respiration.

## Materials & Methods

### Biological model and rearing condition

The Japanese medaka fish (*Oryzias latipes)* is a small, amphidromous freshwater fish from Southeast Asia. This species is eurythermic with a thermal range from zero to 40 °C (Dhillon and Fox, 2007; Hirshfield, 1980; Shima and Mitani, 2004; Kirchen and West 1976), and a thermal optimum of 25°C (Kirchen and West, 1976; Hirshfield, 1980; Leaf et al., 2011). It has a short generation time of 3 months and can reach sexual maturity in about 10 weeks (Hirshfield, 1980), making it a good biological model for multigenerational thermal experimentation. In its native range, the medaka generally lives in small bodies of water such as rice paddies and with important temperature fluctuations (Hirshfield, 1980). In shallow environments, adult medaka can support temperature as high as 43°C (personal observation).

Medaka used in this study were reared over multiple generations at either 20°C (COLD fish) or 30°C (WARM fish) under controlled experimental conditions in climate-controlled chambers. Two lineages by temperatures (W1, W2 for WARM fish and C1, C2 for COLD fish) were derived from an initial F_0_ population of 320 Medakas (with an equal sex ratio) belonging to the CAB strain supplied by Carolina Biological Supply Company (USA) and provided by AMAGEN© (France). The F_0_ generation was initially maintained at 25°C before being gradually acclimatized to 20 or 30 °C (one degree every 2 days). To establish each new generation, eggs clutches were collected during the optimal fecundity period and incubated at the parental temperature (20 °C or 30 °C). We kept the new-born fish in climate-controlled chambers and then housed in five replicates aquariums (25 × 40 × 20 cm) until reaching maturity, with approximately 30 adult fish per tank. They were fed “*ad libitum*” with TétraMin© dry flakes three times daily.

### Whole organism thermal critical maximum tolerance (CT max)

After 5 years of multigenerational exposure to 20°C and 30°C, we exposed fish from each thermal origin to a thermal tolerance dynamic test to estimate their CT_max_. We used 32 fish from the two different thermal origin (W1 & W2, C1 & C2). Specifically, we used 16 fish maintained at 20°C for six generations (F_6_): 8 adults females (4 from C1 & 4 from C2) (mean age in days ± se: 215.1 ± 6.7; i.e. 4.302 degree days) and 8 adults males (4 from C1 & 4 from C2) (215.1 ± 6.7). We also used 16 fish maintained at 30°C for ten generation (F_10_): 8 adults females (4 from W1 & 4 from W2) (122.6 ± 3.8 days; i.e. 3.678 degree days) and 8 adults males (4 from W1 & 4 from W2) (122.6 ± 3.8 days). Ages of the studied fish were chosen accordingly to female reproductive optimum in terms of clutch size (i.e. number of eggs laid per day and per female; Loisel, 2019). Although fish had different ages in days, their physiological ages in degree-days were relatively close (i.e. 3.678 degree days for WARM fish and 4.302 degree days for COLD fish).

WARM and COLD fish used in the experiments present differences in generation numbers (10 and 6, respectively) due to a shorter generation time at 30°C compared to 20°C. Differences in the number of generation is due to warming condition applied in this study, as fish grow faster and have an early sexual maturity when temperature increase, resulting in asynchronous generations between the two acclimation temperatures (Alberto-Payet et al., 2022; Atkinson, 1995, 1994). To get the same fish generation, the experiment should have been run for additional years. Therefore, we preferred comparing mitochondrial respiration of fish at the same date rather than at the same generation to reduce the risk of changes in experimental conditions experienced by WARM and COLD fish (e.g. water quality or potential diseases) over a year, as well as potential changes in conditions of oxygen flux measurements. A previous study on the medaka fish (Loisel, 2019) testing the impact of warm multigenerational exposure, showed that there was few differences in fish growth and reproduction pattern between 20°C and 30°C over 10 generations of exposure. In addition, a common garden experiment revealed that these differences were mainly due to phenotypic plasticity and not to genetic evolution (Loisel, 2019).

In addition, to test if the CT_max_ could influence mitochondrial respiration, 16 control fish (i.e. 4 males and 4 females from each thermal origin) were also maintained under the same laboratory conditions except that they were not exposed to the CT_max_ test.

To estimate the CT_max_, two water baths (WB) were used (WB1 and WB2) (see appendix Fig. S2). Each WB had a water pump and a water heater (1500 Watt). Within each WB, two aquariums were placed at each side of the WB (hereafter referred as to top and bottom positions T & B; Fig. S2). To reduce bias, the location of the fish in the aquariums (i.e. T or B) and in the WB (i.e. WB1 & WB2) was switched each day. The water for the experimental tanks was filtered using two mechanical filters (25 µm then 5 µm) and a UV filter to eliminate microorganisms. Two cameras were installed at the front and at the top of the water-baths to observe fish behaviour remotely. Before the CT_max_ experiment, each fish was weighed, measured, and photographed.

For each thermal origin and lineage, pairs (one male, one female) were placed in an aquarium in the WB at their rearing temperature (20°C or 30°C). Control fish were placed in a recovery aquarium. We then exposed organisms to rapid temperature changes through a constant heating rate (∼ 1.5°C every 10 minutes) that allows deep body temperature to follow ambient temperatures without a significant time lag (Hutchison 1961, 1976). Loss of equilibrium (LOE) was used as an end-point for this experiment (Ziegeweid et al., 2008). Fish were then immediately returned to a tank at their initial temperature to allow their total recovery. Temperature was recorded continuously during the CT_max_ trial. Another potential factor of metabolic stress is a lower oxygen concentration in warmer waters, while metabolic demand increases with temperature. To verify that the oxygen concentration was not limiting, we mesured dissolved oxygen concentration before and after each temperature rise in each aquarium using a fiber optic oxygen probe (*HACH - HQ40d*). For the recovery phase, fish were fed twice a day and monitored individually to ensure their recovery (normal behaviour, weight gain, absence of lesions) over the following 20 days (see appendix Tab. s1).

### Measurement of mitochondrial oxygen consumption rates (OCRs)

After 20 days of recovery following the CT_max_ experiment, we measured their mitochondrial OCR at 25 °C (Topt) and at mean CT_max_ for each thermal origin (WARM and COLD), defined as 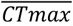, to determine the effects of transgenerational long-term temperature on mitochondrial functioning. We used a full factorial experimental design (see appendix Fig. S3A,B) with (I) fish from the two ancestral temperatures (COLD or WARM) tested at (II) three assay temperatures: 25 °C (thermal optimum), 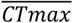 of COLD fish and 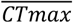 of WARM fish (obtained from the previous experiment). The 25°C assay temperature was chosen to compare fish from both lines at a common non-stressful temperature representing the thermal optimum.

Medakas fish were euthanized by cranial percussion and subsequently measured (mm), weighed (mg), and dissected to collect cerebral tissue samples. Dissections were performed within five minutes post-euthanasia at 4 °C. For each fish, the brain was transferred in 0.5M Respiration Medium (MIR) (Pesta and Gnaiger, 2012), gently mechanically permeabilized with fine forceps and placed on an orbital shaker at 4 °C for 5 min. The permeabilized tissue was split into three different parts and weighed before measurements. Mitochondrial oxygen consumption rates (OCRs) (pmol O_2_·s^-1^·mg^-1^ of tissue) were measured using three Oxygraph-2k high-resolution respirometers (Oroboros Instruments, Innsbruck, Austria), each equipped with two independent measurement chambers. For each fish, we tested simultaneously three assay temperatures (25 °C, 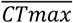 COLD fish, 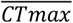 WARM fish) with three Oroboros Oxygraph. To avoid pseudo-replication issues, we switched the temperature of the three Oroboros every day. The oxygen electrodes were calibrated at air-saturated MiR05 and zero oxygen following sodium dithionite addition for each tested temperature. OCR was determined using a sequential substrate/inhibitor protocol (see Appendix Fig. S4).

Specifically, the permeabilized cerebral tissue sample was added to the oxygraph chamber and we immediately injected 5mM Pyruvate, 2mM Malate and 5mM glutamate. These substrates allowed to determine the LEAK value for proton gradient created by electron transfer in complex I of the electron transport system, which represents the oxygen consumption compensating for the proton leak when ATP synthase is not active (non-phosphorylating state). Then, the oxidative phosphorylation at the level of complex I (OXPHOS-CI) was initiated by adding a 2.5 mM saturating concentration of ADP to couple the proton gradient created by electron transfer in CI to phosphorylation of ADP to ATP. Additional injection of cytochrome C (10µM) was used to verify the outer mitochondrial membrane integrity (CYTO-C). The oxidative phosphorylation at the level of complex I+II (OXPHOS-CI+II) was measured with the addition of 10mM succinate. Electron transport system capacity (ETS-CI+II) (reflecting the maximal mitochondrial respiration induced in an uncoupled state) was achieved with repeated additions of carbonyl cyanide p-(trifluoromethoxy) phenylhydrazone 0.5µM (FCCP, 1 + 1 + 1 µL). This was followed by inhibition of complexes I and III through the sequential addition of 0.5µL rotenone and 2.5µM Antimycin A, respectively. The resulting residual oxygen consumption or non-mitochondrial oxygen consumption obtained was subtracted from the previous rates obtained to obtain only mitochondrial specific OCRs. The COX-activity flux was then reached by adding 2mM of Ascorbate and a stepwise addition (5 μL + 5 μL + …) of 0.5mM TMPD. Finally, we added (≥100mM) sodium azide to inhibit the complex IV and correct for auto-oxidation of TMPD. Oxygen flux stabilizes within a few minutes following each injection. The average flux value was retrieved using Datlab software (Oroboros Instruments) to obtain OCRs. We obtain 5 OCRs (LEAK-CI, OXPHOS-CI, OXPHOS-CI+II, ETS-CI+II and COX) each representing distinct processes within the mitochondrial respiratory chain (Fig. S1; Fig.S4).

### Mitochondrial Ratios

The Respiratory Control Ratio of complex I (RCR.CI) which reflects the coupling between electron transport and oxidative phosphorylation from substrates oxidised at complex I, was calculated as the ratio of ATP-synthesizing mitochondrial respiration (OXPHOS-CI) over mitochondrial respiration in the absence of ATP-synthesizing (LEAK-CI) with complex I-induced respiratory substrates (Brand and Nicholls, 2011; Estabrook, 1967). The respiratory control ratio represent a single general measure of mitochondrial function. In permeabilized fish brain (as used in this study) the RCR with complex I substrates (RCR.CI, Eq.1) has been shown to fall into the range of 1.4 to 7.5 during control conditions depending on species and thermal origins (Chung et al., 2017; Devaux et al., 2019; Devaux et al., 2023).

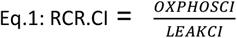

We also calculated the substrate relative contribution ratio (SCR, Eq.2) (i.e. the relative contribution to increased oxygen consumption rate when adding a new substrate) of complex I + II (succinate) to complex I only (Cytochrome-C) for OXPHOS state:

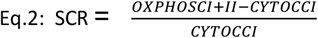

Finally, for each fish, we calculated ratios of OXPHOS-CI, LEAK-CI and OXPHOS-CI+II values measured at their 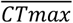 divided by 25 °C assay temperature values (i.e.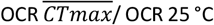) to characterize the fold change in mitochondrial oxygen consumption at the CT_max_ relative to the oxygen consumption at the optimal temperature.

### Statistical analyses

Statistical analyses were performed using R software version 4.3.1 (R Foundation for Statistical Computing Platform, Free Software Foundation’s, Boston, USA). For the CT_max_ experiment, we used a linear mixed effect model (LMM) to test the effects of fish ancestral temperatures (i.e. 20 °C or 30 °C), fish lineage, fish sex and their interactions on the 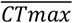 values. The aquarium position nested in the experimental day (i.e. the same information as WB) were included as random effects, to account for potential spatial and temporal effects. Non-significant terms were excluded from the final analysis.

For OCRs at the mitochondrial level, we tested the effects of fish ancestral temperatures, assay temperatures, treatment (CT_max_ test or control fish), fish sex, fish lineage, and their interactions for LEAK-CI, OXPHOS-CI, OXPHOS-CI+II, ETS-CI+II and COX activity as well as on the RCR.CI and SCR using LMMs. The fish identification number and the couple pair code nested in the Date experiment were included as random factors, to account for the repeated measurements (three measurements for each fish) and for potential temporal effects. Non-significant terms were excluded from the final analysis. For each model, when the interaction was significant between the two experimental conditions tested, we divided the datasets by the experimental condition using LMMs. OCRs presenting anomaly during measurements or not having reacted to injections were excluded from the analyses.

We then compared OCRs between the COLD fish at their assay temperate 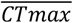 (C-38.9) and the WARM fish at their assay temperature 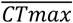 (W-43.8) using LMMs to better understand if their mitochondria function similarly when exposed to their respective CT_max_ temperatures.

We finally illustrated distribution pattern of the individual CT_max_ for each fish for OXPHOS-CI, LEAK-CI and OXPHOS-CI+II as a function of the mitochondrial OCRs at 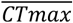 assay temperature. We also compared distribution pattern for the OCR at 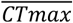 divided by the OCR at 25°C for OXPHOS CI, LEAK CI & OXPHOS CI+II as a function of CT_max_ whole-organism values for each thermal origin and tested the differences in the Ratio (flux CTM/ flux 25°C) between the two fish thermal origins using LMMs.

Application conditions for the LMMs were tested. The significance of the fixed model terms was assessed using analyses of deviance (Anova function from the car package) and non-significant (p >0.05) variables or interactions were removed from the final model. Linear mixed models (LMMs) were implemented using the lme4 package (Bates et al., 2015).

## Results

### Whole organism thermal critical maximum (CT_max_)

The 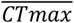 values (Fig. 1) were significantly influenced by fish ancestral temperature (Chisq = 163, df = 1, p < 0.001 ***, Fig. 1; see appendix Tab. S2). Specifically, WARM fish had a 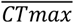 (43.8 °C ± 0.27) significantly higher than COLD fish (38.9 °C ± 0.32) representing a 12.6% increase.

**Figure 1:**
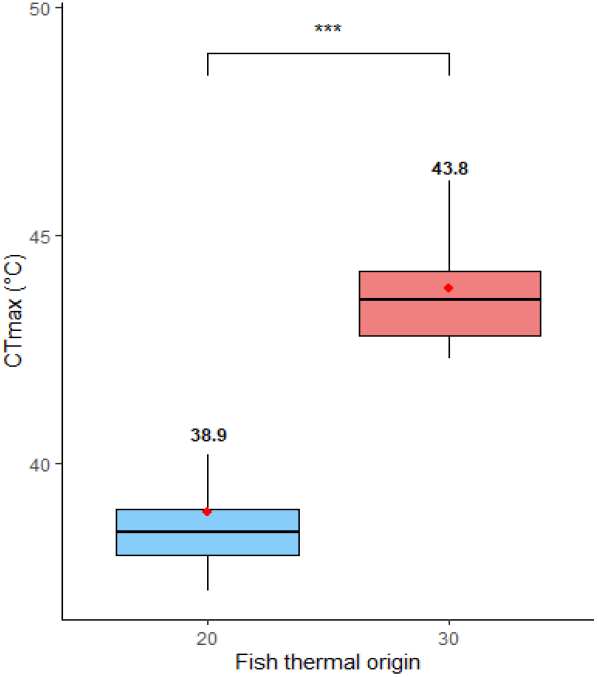
Boxplot with Mean (red dots) of the CT_max_ (°C) according to fish ancestral temperature (20 for COLD fish & 30 for WARM fish) (N= 32). *** represents highly significant differences between WARM and COLD fish (P < 0.001).

### Mitochondrial oxygen consumption rates (OCRs)

LEAK-CI (Fig. 2a; see appendix Tab. S3, S4) was not influenced by the interaction between ancestral temperatures and assay temperatures (Chisq = 4.9, df = 2, p = 0.09). However, LEAK-CI OCR was significantly influenced by assay temperature (Chisq =16.5, df =2, p < 0.001 ***) through increased OCR with temperatures. We found no significant effect of ancestral temperatures on LEAK-CI (Chisq = 1.05, df = 1, p = 0.31).

**Figure 2:**
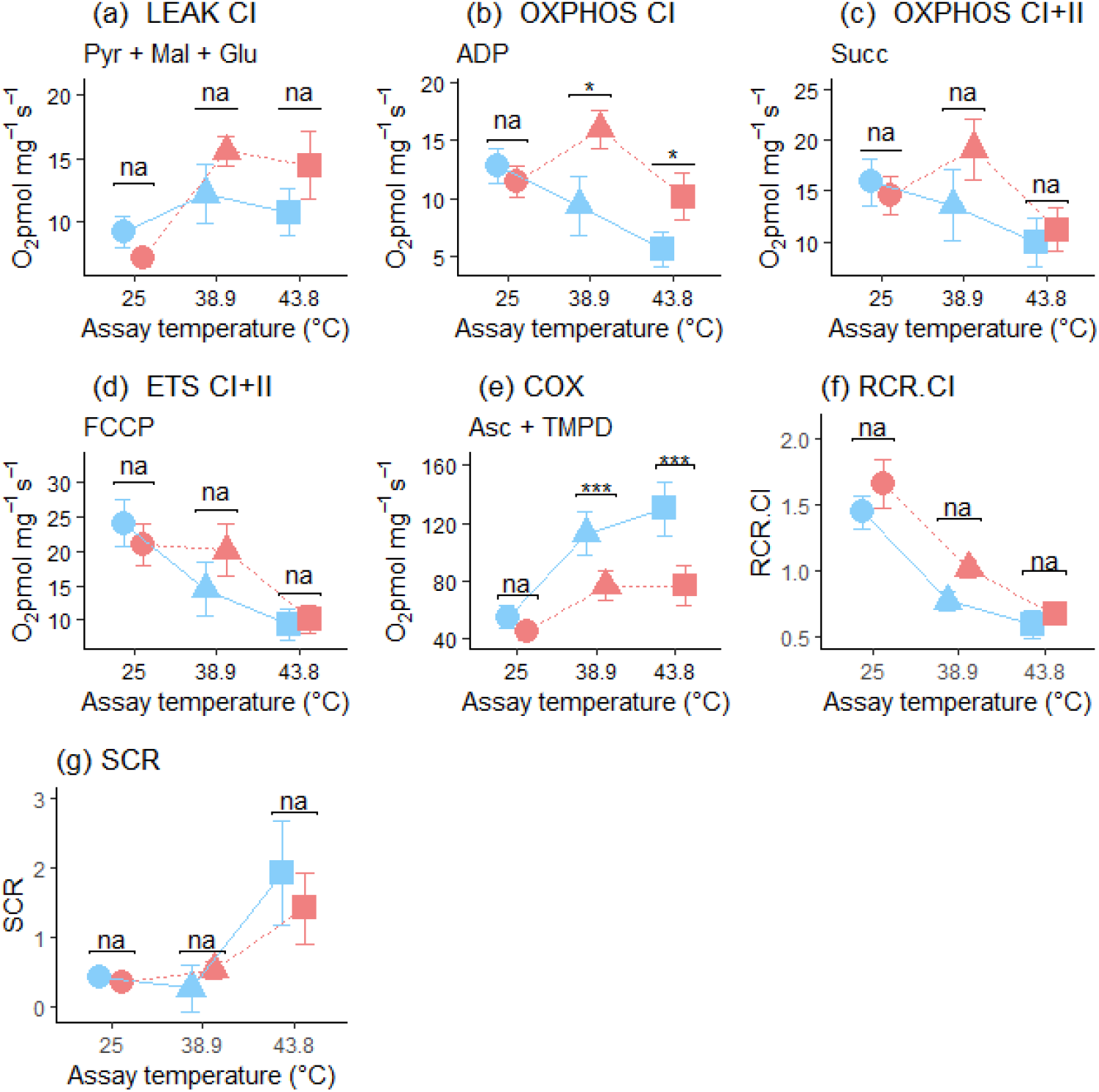
(a) LEAK-CI, (b) OXPHOS-CI, (c) OXPHOS-CI+II, (d) ETS-CI+II, (e) COX, (f) RCR.CI and (g) SCR mean (± SE) oxygen consumption rates (OCRs) in picomol.s^-1^ .mg^-1^ according to the ancestral temperature (COLD 20 °C fish in blue or WARM 30 °C fish in red) and assay temperature (25 °C, dots; 38.9 °C, triangles; 43.8 °C, square). * represents significant differences between WARM and COLD fish (P < 0.05).

OXPHOS-CI (Fig. 2b; Tab. S3, S4) was significantly influenced by the interaction between ancestral temperature and assay temperature (Chisq = 9.94, df = 2, p = 0.006 **). When we split the dataset by experimental conditions, COLD and WARM fish OXPHOS-CI state was significantly affected by assay temperature. For COLD fish, we observed a decrease with increased assay temperature (Chisq = 12.39, df = 2, p < 0.005**) and for WARM fish we observe an increase between 25 °C and 38.9 °C followed by a decrease between 38.9 °C and 43.8 °C (Chisq = 12.16, df = 2, p < 0.005**). Moreover, we found a significant difference between COLD and WARM fish at the 38.9 °C assay temperature (Chisq = 4.8, df = 1, p < 0.05*) and at 43.8 °C (Chisq = 4.21, df = 1, p < 0.05*) but no difference for 25 °C (Chisq = 0.3, df = 1, p =0.59).

OXPHOS-CI+II was significantly influenced by the interaction between ancestral temperature and assay temperature (Chisq = 6.14, df = 2, p =0.05*) (Fig. 2c; Tab. S3, S4). When we split the dataset by experimental conditions, OXPHOS-CI+CII was significantly influenced by assay temperature for COLD and WARM fish. For COLD fish, we observed a decrease with increased assay temperature (Chisq = 6, df = 2, p =0.05*), and for WARM fish, an increase between 25 °C and 38.9 °C followed by a decrease between 38.9 °C and 43.8 °C (Chisq = 21.3, df = 2, p < 0.001***). We found no significant difference between COLD and WARM fish for each assay temperature (25 °C: Chisq = 1e-04, df = 2, p =0.99; 38.9 °C: Chisq = 1.47, df = 2, p =0.23; 43.8 °C: Chisq = 0.42, df = 2, p =0.52).

ETS-CI+II (Tab. S3, S4; Fig. 2d) was not significantly influenced by the interaction between ancestral temperature and assay temperature (Chisq = 5.93, df = 2, p = 0.051) even if we are close to the interaction and we are seeing similar trends to the OXPHOS fluxes. ETS-CI+II significantly decreased with assay temperature (Chisq =46.33, df =2, p = p < 0.001 ***). Finally, we found no significant effect of ancestral temperature on ETS-CI+II (Chisq = 0.36, df = 1, p = 0.55).

The COX-activity (Tab. S3, S4; Fig. 2e) was not significantly influenced by the interaction between ancestral temperature and assay temperature (Chisq =3.48, df = 2, p =0.18). This OCR significantly increased with assay temperature (Chisq =23.4, df =2, p = p < 0.001 ***). We also found a significant effect of ancestral temperature (Chisq = 11.5, df = 1, p < 0.001 ***).

For all OCRs, we observed a non-significant general trend towards lower fluxes for WARM fish at 25 °C than for COLD fish at 25 °C.

### Mitochondrial Ratio

The RCR.CI was generally low for all conditions tested (Tab. S3; Fig. 2f). It is however important to note that RCR measured using complex I substrates in permeabilized fish brain is highly dependent of species and has been reported to be as low as 1.4 during control conditions (Devaux et al., 2019; Devaux et al., 2023). The RCR.CI measured here is also only taking into account complex I substrates which might not be optimal oxidative substrates to fuel mitochondrial respiration in medaka fish. The RCR.CI was not significantly affected by the sex and the treatment, but we observed a significant effect of lineage (see appendix Tab. S5). We found the same tendency for the two lineages through lower fluxes for COLD fish than WARM fish. This tendency was non-significant for C1 and W1 but (W2) and COLD (C2) displayed significant differences at 25 °C (Chisq = 7.7, df = 2, p = 0.005**) and 38.9 °C (Chisq = 5.2, df = 2, p = 0.03*) (see appendix Fig. S5). RCR.CI was not significantly influenced by the interaction between ancestral temperature and assay temperature (Chisq = 0.95, df = 2, p = 0.62), but was significantly influenced by assay temperature (Chisq =94.2, df =2, p < 0.001 ***) through a decrease with increased assay temperature. We found no significant effect of ancestral temperature on the RCR.CI (Chisq = 1.76, df = 1, p = 0.18).

The statistical interaction between ancestral temperature and assay temperature was non-significant (Chisq = 1.13, df = 2, p = 0.57) for the SCR ratio reflecting the succinate relative contribution to respiration (Tab. S2, S5; Fig. 2g). However, the SCR was significantly influenced by assay temperature (Chisq =15.5, df =2, p < 0.001 ***) through an increase with assay temperature. We found no significant effect of ancestral temperature on SCR (Chisq = 0.11, df = 1, p = 0.73).

### OCRs comparisons between warm and cold fish at their respective CT_max_

To highlight mitochondrial OCRs differences at 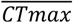 temperature between thermal lineages, we compared OCRs between COLD fish at 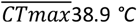 and WARM fish at 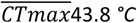 (Tab. S3; Fig. 3). We found that OCRs did not significantly differ between COLD and WARM fish at their respective 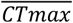 for LEAK-CI (Fig. 3a) (Chisq = 0.91, df = 1, p = 0.34), OXPHOS-CI (Fig. 3b) (Chisq = 0.79, df = 1, p = 0.37), OXPHOS-CI+II (Fig. 3c) (Chisq = 0.25, df = 1, p = 0.62), ETS-CI+II (Fig. 3d) (Chisq = 1.17, df = 1, p = 0.28), RCR.CI (Fig. 3f) (Chisq = 3.49, df = 1, p = 0.38) and for SCR despite slightly higher SCR for WARM fish (Fig. 3g) (Chisq =3.5, df =1, p = 0.06). However, we found that COX activity significantly differed between COLD and WARM fish at their 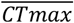 (Fig. 3e) (Chisq = 4.38, df = 1, p = 0.03*).

**Figure 3:**
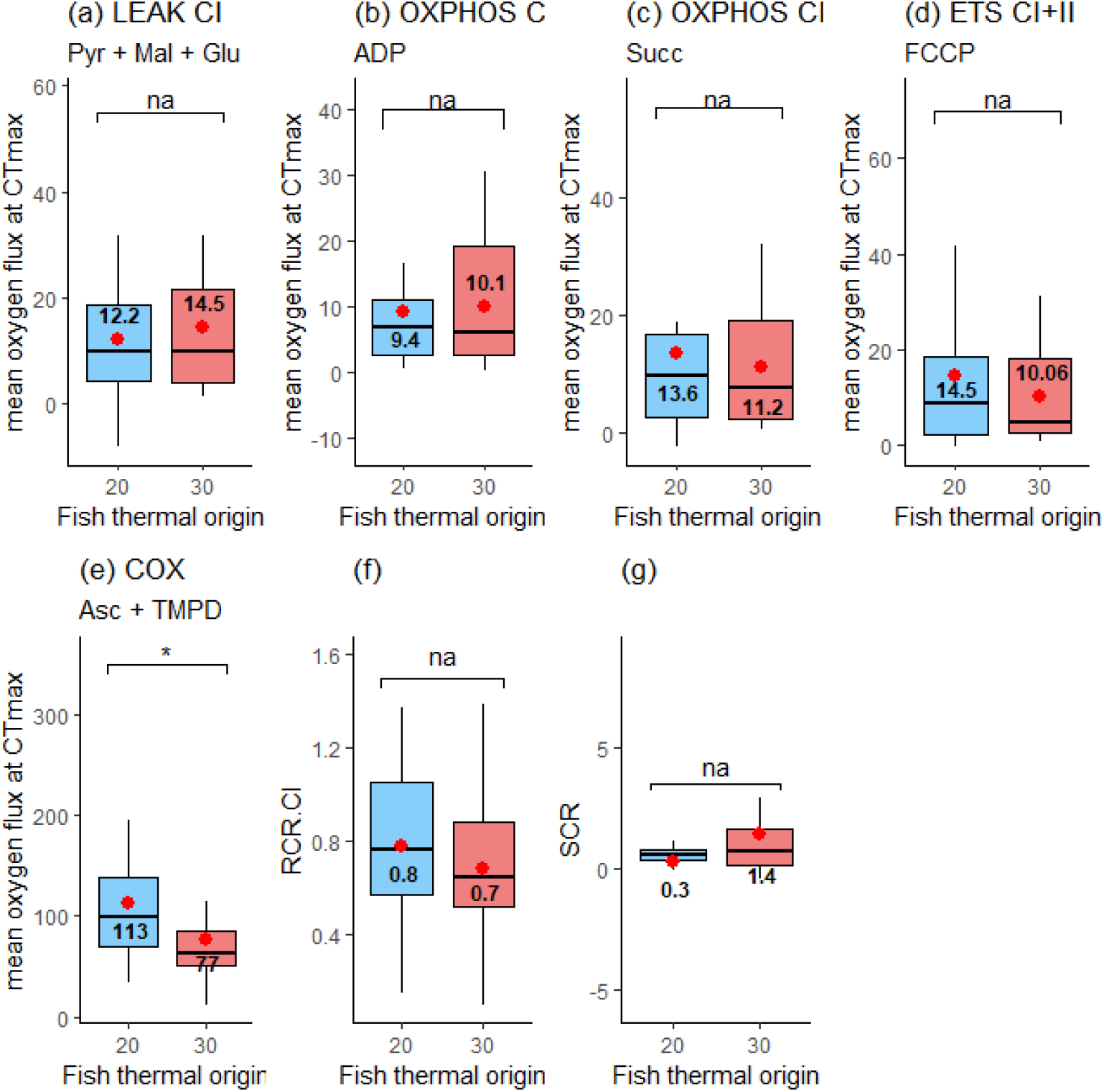
Boxplot with mean (red dots with black text) of OCR in picomol.s^-1^ .mg^-1^ at 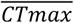 assay temperature for (a) LEAK-CI, (b) OXPHOS-CI,(c) OXPHOS-CI+II,(d) ETS-CI+II, (e) COX activity, (f) RCR.CI, and (g) SCR for each thermal origin (COLD in blue or WARM in red). * represents significant differences between WARM and COLD fish (P < 0.05).

### Links between mitochondrial OCRs and whole organism CT_max_

For a better understanding of the link between whole organism CT_max_ and mitochondrial OCRs at 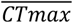 assay temperature, we plotted the 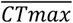 assay temperature according to the whole-organism real CT_max_ values for each fish against individual mitochondrial OCRs *in picomol*.*s*^*-1*^ .*mg*^*-1*^ (see appendix Fig. S6) as well as for relative OCRs (i.e. OCR 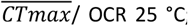; see appendix Fig. S7)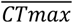. We did not observe a distribution pattern of OCR or ratio values flux at 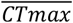 assay temperature for OXPHOS-CI, LEAK-CI and OXPHOS-CI+II (Fig. S6 and Fig. S7). We also illustrated the fold change in mitochondrial OCR at 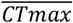 temperature relative to the OCR at the optimal temperature 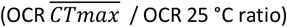 for LEAK CI, OXPHOS CI & OXPHOS CI+II according to the whole-organism real CT_max_ values for each fish from both thermal origins (Fig. 4). We found no significant difference in the Ratio 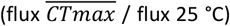 between the two fish thermal origins for LEAK-CI (Chisq = 0.02, df = 1, p = 0.9), OXPHOS-CI (Chisq = 0.10, df = 1, p = 0.74) and OXPHOS-CI+II (Chisq = 1.12, df = 1, p = 0.29) (Fig.4).

**Figure 4:**
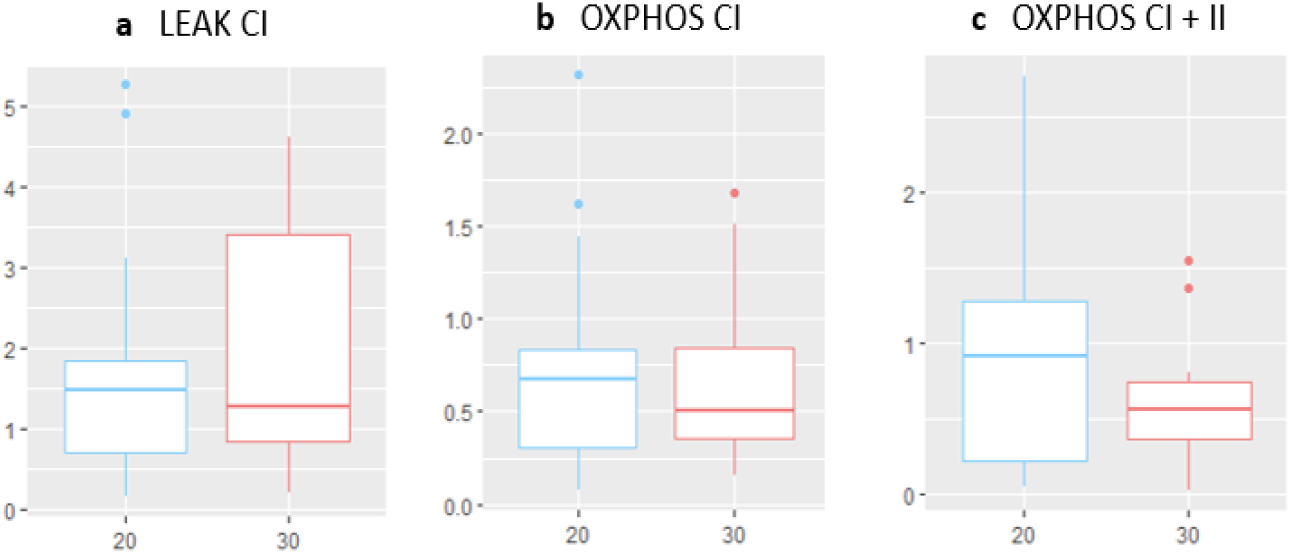
Boxplot of the Ratio (OCR 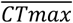/ OCR 25 °C) for (a) LEAK-CI, (b) OXPHOS-CI and (c) OXPHOS CI+II, according to fish thermal origin (i.e. ancestral temperature of 20 °C for COLD fish in blue or 30 °C for WARM fish in red).

## Discussion

The aim of this study was to investigate whether acclimation over several generations (multigenerational exposure) of freshwater Medaka fish to an elevated, non-stressful temperature (30°C) can result in changes in whole-organism thermal limits (CT_max_) and brain mitochondrial functions. Moreover, we aimed at determining whether there is a link between whole-organism CT_max_ and brain mitochondrial performance at elevated temperature in the same fish. These questions are motivated by studies showing that higher thermal tolerance can be improved after a period of exposure to elevated temperature (Bilyk and DeVries, 2011; Angiletta 2001) and by studies showing that mitochondrial neural dysfunction can limit the upper thermal tolerance of the entire organism (Somero and DeVries, 1967; Ern et al., 2015).

### Whole organism upper thermal tolerance (CT_max_)

In line with studies suggesting that the CT_max_ varies with the exposure duration (Podrabsky and Somero, 2006; Bilyk and DeVries, 2011; Angilletta 2001), we found that thermal acclimation over several generations increases the maximum thermal tolerance of the whole organism. Fish from multigenerational exposure at 20 °C (COLD fish) had a 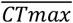 of 38.9 °C, compared to 43.8 °C for those exposed to 30 °C (WARM fish) representing a 12.6% increase (Fig.1).

Our results thus highlight the influence of an acclimation period spanning several generations (long-term exposure) on the thermal tolerance of the whole organism. Similar results have already been observed for studies performing single-generation exposure. For example, Fangue et al., (2006) obtained significant differences in CT_max_ for common killifish (*Fundulus heteroclitus*) after 21 days of acclimation. Killifish acclimated at 5 °C had a CT_max_ of 31 °C, while fish acclimated at 33 °C had an average CT_max_ around 41 °C (representing a 10°C increase). Li et al., 2015 also demonstrated that the CT_max_ of marine medaka (*Oryzias melastigma*) larvae increases significantly with acclimation temperatures. Fish were subjected to stepwise temperature change at a rate of 1 °C/h, increasing or decreasing from a starting temperature of 25 °C (the control) to reach six target temperatures (12, 13, 15, 20, 28 and 32 °C) with a range of maximum thermal tolerance variation from 39.9 °C at 13 °C to 42.8 °C at 32 °C. In their study, they found a CT_max_ of 40.53 °C at 20 °C and a CT_max_ of 42.56 °C at 30 °C (Li et al., 2015). Since exposition took place over several generations in our case, it is even possible that this difference in thermal tolerance reflects adaptation to different thermal regimes considering that temperature might be a driving force for species adaptation. Genetic and genomic data would however be essential to evaluate the relative contribution of phenotypic plasticity (acclimation) versus genetic adaptation for the observed changes in CT_max_.

### Mitochondrial performances

The link between acute warming and mitochondrial function is already well established. Several studies have shown that acute exposure to elevated (non-stressful) temperatures leads to an increase in mitochondrial respiration (Abele et al., 2002; Bielski et al., 1983; Brand, 2000; Ricquier and Bouillaud, 1998; Roussel and Voituron, 2020). However, this increase in mitochondrial respiration can also lead to higher levels of ROS (Reactive oxygen species) production at temperatures close to the thermal limit that can fasten ectotherms senescence and mortality (Christen et al., 2018; Woodward, 2009). In this study, we observed a general increase in OCRs for OXPHOS-CI, OXPHOS-CI+II and LEAK-CI states between assay temperatures of 25 °C and 38 °C for WARM fish (38°C is considered as non-stressful because lower than 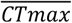 value for WARM fish), confirming that assay elevated temperature exposure leads to an increase in oxygen consumption (Fig. 2abcd).

There is evidence of thermal mitochondrial plasticity in ectotherms following acclimation (Guderley and St-Pierre, 2002; Seebacher et al., 2010, Morla et al., 2024). Our results indicate a trend towards lower fluxes for WARM fish at their optimal temperature (25°C assay temperature). Similar results have been observed previously in other ectotherms (Chung et al., 2017; Ekström et al., 2016; Khan et al., 2014; Schulte, 2015; Yan and Xie, 2015) and on medaka lineage after two years of temperature exposure (Morla et al., 2024). This decline in mitochondrial respiration with acclimation would allow counteracting the increasing oxygen consumption inherent to faster reaction rates at higher temperature and associated cellular damages (Abele et al., 2002; Hochachka and Somero, 2002).

Interestingly, regarding LEAK-CI, mitochondrial fluxes increased between assay temperatures of 25 °C and 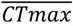, regardless of the thermal origin of fish (Fig. 2a). Previous studies have reported mechanisms allowing to reduce the metabolic cost at elevated temperatures by decreasing mitochondrial efficiency in term of ATP production through increasing proton leakage (LEAK) and decreasing the proton gradient for ATP production (Brand, 2000; Ricquier and Bouillaud, 1998; Skulachev, 1998). In our case, RCR.CI values were very low (< 1) (Fig.2f) for assay temperatures above 25°C. Studies on fish brain mitochondrial functions usually report low to very low oxygen fluxes (and thus RCR) during control conditions with complex I substrates using either isolated mitochondria or permeabilized brain tissue (Cambier et al., 2012; Bourdineaud et al., 2013; Chung et al., 2017; Devaux et al., 2019; Willis et al., 2021). In the context of acute thermal challenge, these fluxes are impaired, leading to even lower RCR indicative of mitochondrial dysfunction (which is also highly species-dependent, Chung et al., 2017; Willis et al., 2021), thus consistent with our RCR.CI values. Moreover, considering that almost no increased oxygen consumption was detected after addition of cytochrome c, which attests of mitochondrial outer membrane integrity, the low values for the RCR.CI may reflect a specific impairment of complex I at high temperature. Alternatively, this indicates that substrates used were not optimal to stimulate oxygen consumption in our species (at least at high temperature) consistent with what has been demonstrated in other freshwater species such as mussels (Hraoui et al., 2020). Nevertheless, our results showing that LEAK flux are higher than OXPHOS flux at high temperature suggest a complete crash of substrate oxidation capacity at complex I relative to what is required to compensate for the proton leak (Brand and Nicholls, 2011). This could result in a mismatch between ATP demand and supply, as impairment of pyruvate and malate oxidation at elevated temperature has been shown to associate with a collapse of ATP synthesis and a consequent drop in the ATP/O efficiency (Roussel et al., 2023). Altogether, these results are strongly indicative of mitochondrial dysfunction and especially at the CI level, which could lead to reduced efficiency in terms of ATP production. However, measuring ATP production would be required to confirm this interpretation.

There was no difference in OXPHOS and LEAK between WARM and COLD fish at 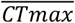 temperature (Fig. 3abcdfg), suggesting a convergence towards a similar response of the mitochondrial OCRs at temperature reaching the 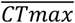. We did not observe neither significant difference in the Ratio 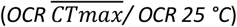 between the two thermal origins of the fish (Fig.4) suggesting limits in thermal plasticity at stressful elevated temperature for mitochondrial respiration.

However, our results concerning the relative contribution of complex II (SCR) (Fig. 2g) show an increase between assay temperatures of 25 °C and 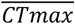 only for WARM fish, suggesting that the relative contribution of complex II at higher stressful temperatures is more important for these fish. This result suggests that WARM fish may exhibit significant mitochondrial flexibility in complex II at temperatures reaching the thermal limit of the organism. A possible explanation for this result would be that WARM fish have developed plastic or genetic strategies to avoid the problem of substrate oxidation capacity at the complex I level (Jørgensen et al., 2021; Menail et al., 2022; Blanchard et al., 2024). This mechanism due to plasticity, genetic and/or epigenetic inheritance (Boukal et al., 2019; Monaghan et al., 2009) could appear after a period of acclimation over several generations at high temperature, in order to compensate for the high thermal sensitivity of complex I identified as the primary site for heat failure in mitochondria (El-Wadawi and Bowler, 1996; Chung et al., 2018; Martinez et al., 2016; Iftikar et al., 2010).

### Links between mitochondrial performances and whole organism CT_max_

Due to the thermal sensitivity of mitochondria, studies have suggested a link between mitochondrial neuronal dysfunction under thermal pressure and the failure of whole-organism (Somero and DeVries, 1967; Pörtner, 2002; Iftikar and Hickey, 2013; Iftikar et al., 2014; Ern et al., 2015). In this study, we did not observe a distribution pattern of ratio or OCR values (Fig. S6 & S7) as a function of relative CT_max_ for each fish measured for LEAK-CI, OXPHOS-CI, and OXPHOS-CI+II.

Our results suggest a decrease (not always significant) in the flux of OXPHOS-CI, OXPHOS-CI+II and ETS-CI+II between assay temperatures of 25 °C and 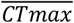, for WARM and COLD fish (Fig. 2 bcd). Chung et al., (2017) found similar results for mitochondrial OCRs of fish acclimatized to 15 °C, which decreased significantly as they approached 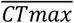, suggesting a link between whole-organism CT_max_ and mitochondrial performance at critical temperature. A more important decrease was observed for WARM fish, suggesting greater thermal sensitivity (Fig. 2bcd). However, the link between whole-organism CT_max_ and mitochondrial performance at elevated temperatures needs to be clarified. Our results also suggest that the mitochondria of COLD fish continue to function (flux > 0) beyond average whole organism mean heat tolerance. In line with this finding, recent studies suggests that mitochondria are more tolerant than the whole organism (Chung and Schulte, 2020; Jørgensen et al., 2021; Syromyatnikov et al., 2019). However, mitochondrial responses to extreme temperatures can also affect other functions, such as ATP production, ROS production, coupling efficiency (ATP/O) and membrane fluidity (Biederman et al., 2019; Roussel et al., 2023) which we have not measured in this study and which could also explain our results.

To conclude, our results indicate that 5 years of multigenerational exposure to elevated temperature significantly increases the CT_max_ of individuals. Although cerebral mitochondria continue to function beyond the average heat tolerance of the whole organism, our results also suggest a link between whole-organism CT_max_ and cerebral mitochondrial performance, particularly in terms of complex I oxidation capacity, highlighting the importance of brain mitochondria in the context of thermal tolerance. Finally, we suggest that WARM fish may exhibit mitochondrial flexibility at the level of complex II at temperatures reaching the thermal limit of the whole organism. Our study highlights the complexity of the relationship between mitochondrial thermal sensitivity and whole-organism thermal tolerance, as well as the fact that acclimation can modify mitochondrial and thermal performance, which can have tremendous impacts on natural populations considering the increased average environmental temperature as well as the frequencies and durations of heatwaves brought forth by climate change. These capacities are nevertheless rarely taken into account in species distribution models (Clusella-Trullas et al., 2021). Finally, thermal tolerances can vary with life stage or body size, we thus need more research to determine if the limiting mechanisms during acute warming also differ across species, life stages, and thermal history (Pörtner et al., 2021; Dahlke et al., 2020).

## Supporting information

Supplementary material for the article (tables and figures)

## Ethical approval

Experimental protocols were approved and performed in agreement with the ethical permits #42422-2023040411317548 v3. The above-mentioned project has been ethically assessed by the ethics committee for animal experimentation n°071.

## Data accessibility

The data that support the findings of this study are openly available on ZENODO at https://doi.org/10.5281/zenodo.14900919

## Acknowledgments & Funding

This work was funded by the project EcoTeBo (ANR-19-CE02-0001-01) from the French National Research Agency (ANR) and by the project CLIMAQUA from the PACA (Provence-Alpes-Côte d’Azur) region and the INRAE AQUA department. We thanks D. Roussel and Y. Voituron for Oroboros equipment.

