## Supplementary material for the article (tables and figures) for "“Multigenerational effects of temperature exposure on upper thermal limit and mitochondrial functioning in Medaka (*Oryzias Latipes*) brain”"

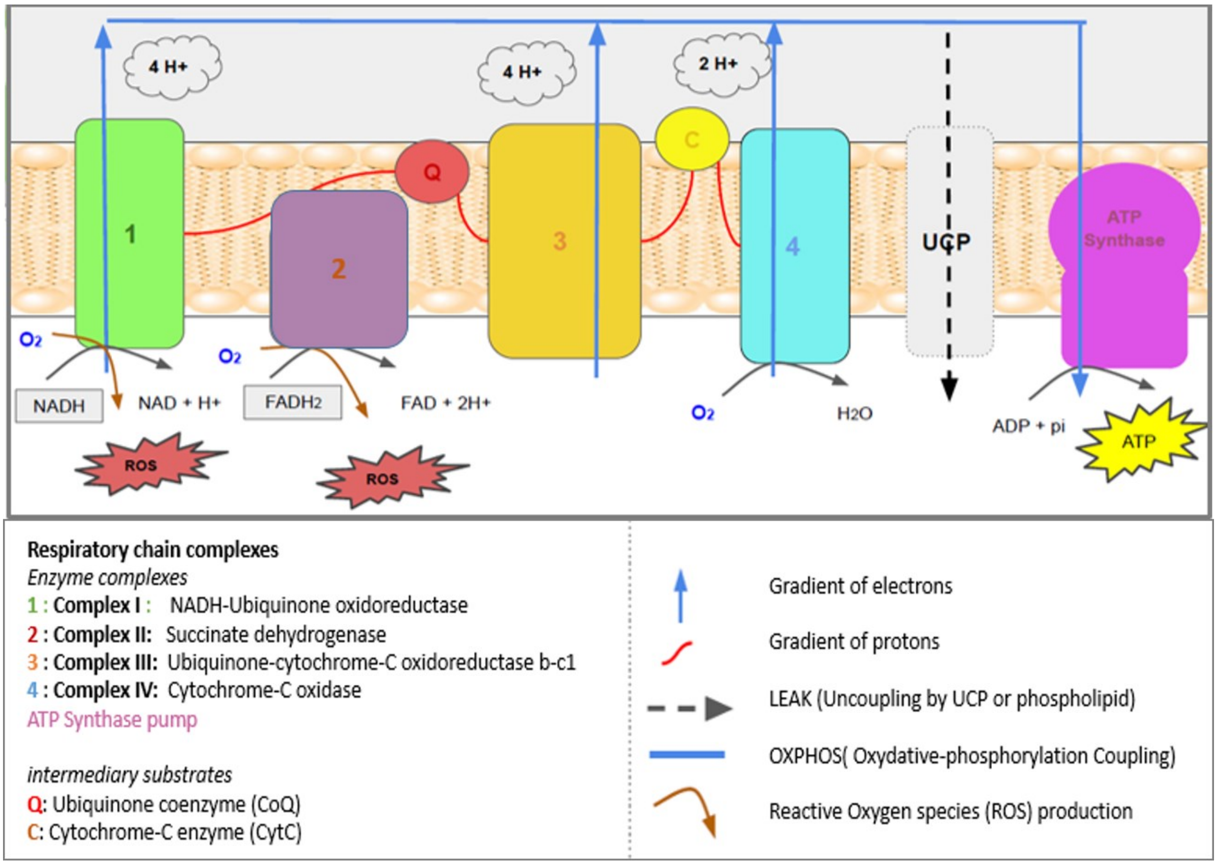


***Fig. S1:*** *The oxidative phosphorylation process in the mitochondria respiratory chain: coupling between cellular respiration and phosphorylation of ADP to ATP. The flow of electrons from NADH and* $FADH_2$ *is guided by the redox potentials of the four complexes and by the coenzymes (Q and C) to the final acceptor, the oxygen which is reduced to* $H_2O$*. Each of these complexes, except 2, couples the flow of electrons with a proton pump. The energy of the proton gradient is used to phosphorylate ADP into ATP. Protons can sometimes return to the matrix independently of ATP synthase, via uncoupling proteins (UCPs) or through the membrane phospholipids (mitochondrial uncoupling). The electron transport leads to the production of ROS, reactive oxygen species.*

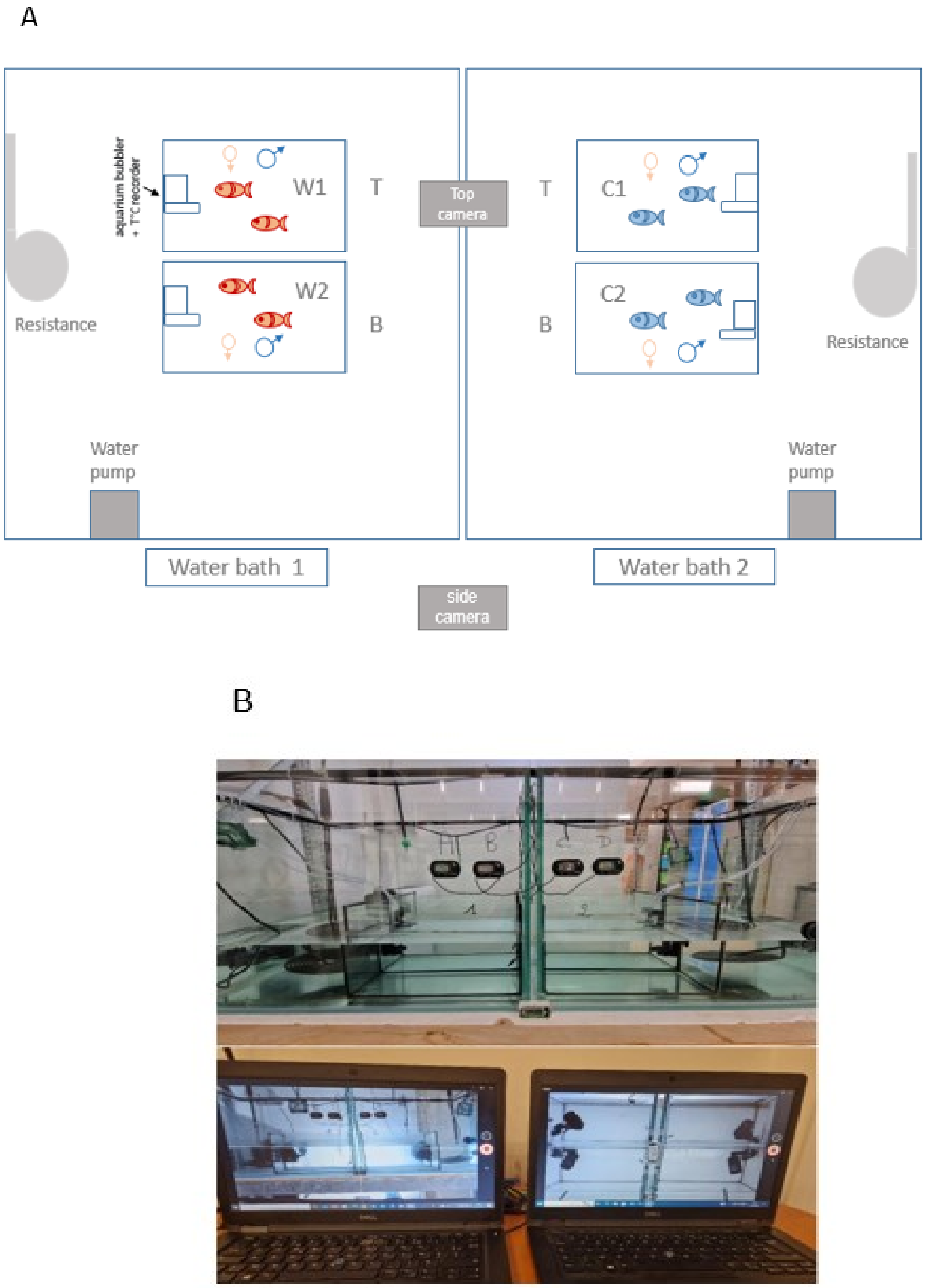


***Fig. S2***: *(A) Schematic top view of the experimental design of the* $CT_{max}$ *experiment with C1 & C2 for COLD lineages and W1 & W2 for WARM lineages; water baths 1 and water baths 2; T or B for top or bottom aquariums. Red fish represent fish reared over several generations at 30°C and blue fish represent fish reared over several generations at 20°C. Fish position was interchanged between aquaria and water baths. (B) Photo of the experimental structure of the water baths and the fish observation device.*

***Tab. S1:*** *health score grid for evaluating clinical condition during 20 day after* $CT_{max}$ *experiment for each fish.*

| Date: | Time | Fish n°.: | Score (from 1 to 4) |
|---|---|---|---|
| Appearance | Body condition | Weight (overweight, skinny, emaciated, cachectic) | |
| | | inflated, e.g. swim bladder, eggs retention | |
| | | Altered shape, e.g. spinal abnormalities | |
| | | Modified or missing fins | |
| | | Modified or missing gill covers | |
| | | Eye lesions | |
| | | Weight loss (5, 10, 15, 20%) | |
| | Scales and skin condition | Scale/skin changes | |
| | | Skin reddening | |
| | | Lighter/darker pigmentation | |
| | | Other skin color changes | |
| | | Ulcers | |
| | | Localized protuberance/tumor | |
| Body function | Breathing | Increased opercular frequency | |
| | | Surface inspiration | |
| | Food intake | Quantity of food consumed | |
| | Other | Please specify | |
| Spontaneous and provoked behaviors | Swimming and balance | Loss of balance, loss of equilibrium | |
| | | Increase/decrease in activity | |
| | | Swimming in circles/corkscrew/spiral | |
| | | Rubbing against aquarium wall | |
| | | Swimming at bottom of aquarium | |
| | | Swimming at surface | |
| | Responses to stimulation | Feeding activity | |
| | | Avoidance of mechanical stimulation | |
| | | Avoidance of light beam | |
| | Social interaction | aggressiveness | |
| Free observation | Body condition | Record observations of unexpected negative effects on well-being here. | |
| | | | Score total : |

(A)

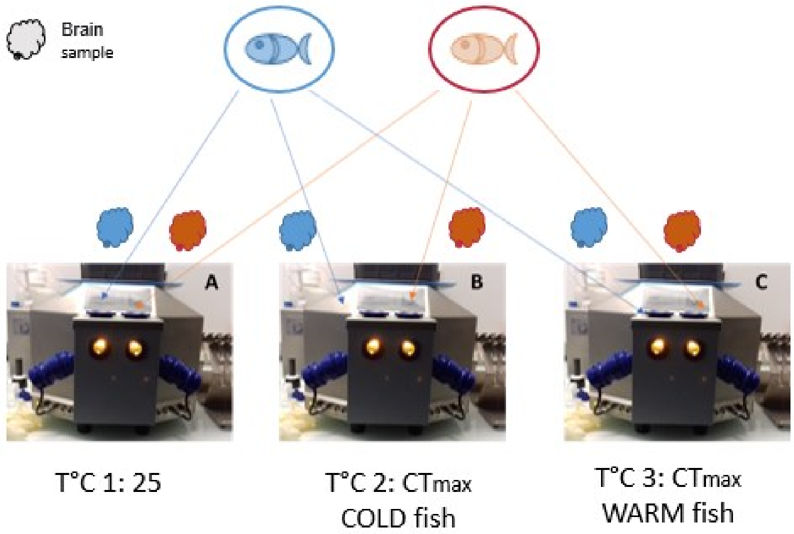


(B)

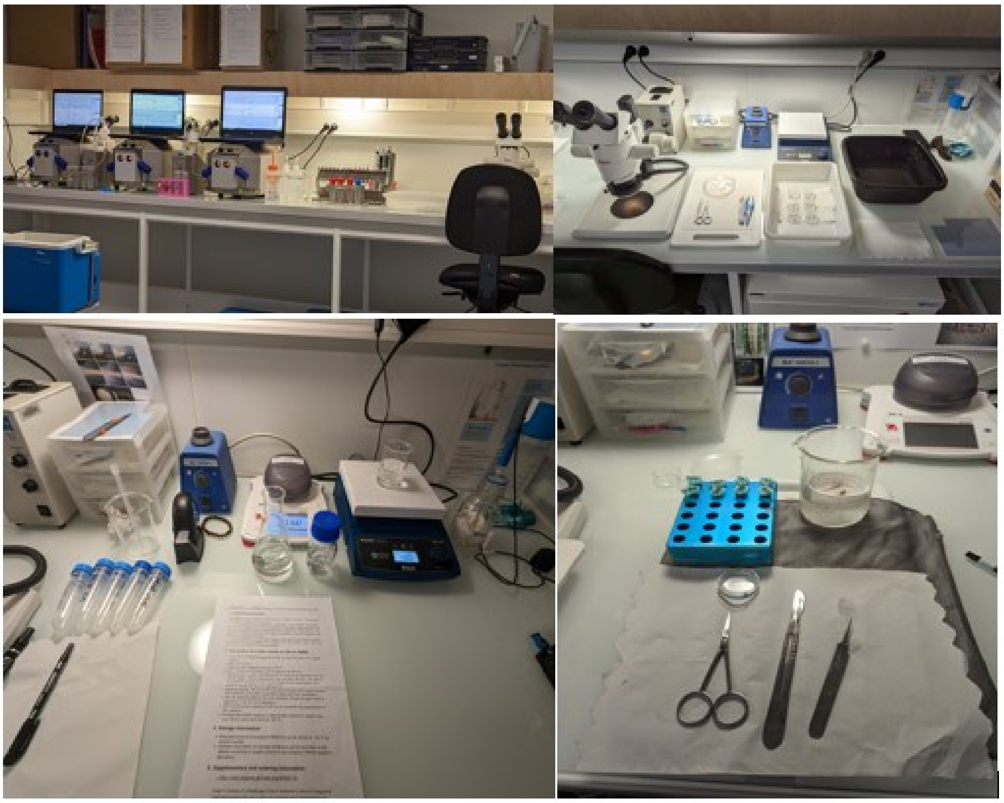

***Fig. S3:*** *(A) full factorial mitochondrial experimental design. Blue fish were reared at 20 °C over several generations (COLD) and red fish were reared at 30 °C over several generations (WARM). A, B, C represent the three oxygraphs with two independent measuring chambers at T°C 1: 25 °C (thermal optimum), T°C 2:* $\overline{CTmax}$ *of fish from 20°C and T°C 3:* $\overline{CTmax}$ *of fish from 30°C. (B) Experimental room and material for dissection and mitochondria respiratory measurement.*

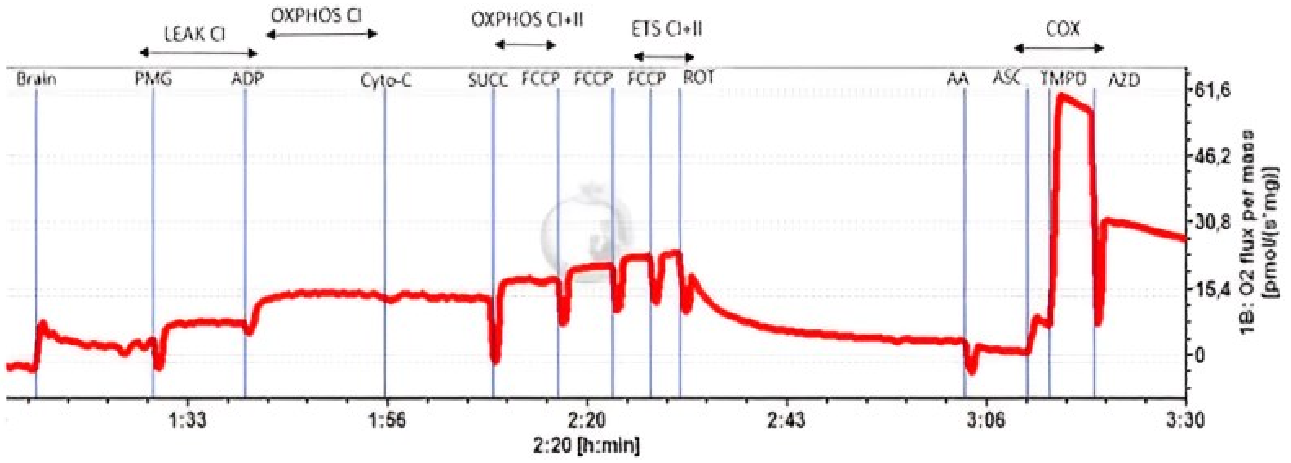


***Fig. S4**: Oxygen flux (picomol.s$^{-1}$.mg$^{-1}$) according to time (h :min) through the sequential activator/inhibitor injection of the respiratory chain.*

**Tab. S2:** *Output of the Analysis of Deviance Table (Type II Wald chisquare tests)* on the most parsimonious models for $\overline{CTmax}$ variable.

| RESPONSE VARIABLE | PREDICTORS (Fixed effets) | DF | CHISQ | *P* |
|---|---|---|---|---|
| $\overline{CTmax}$ | *Lineage* | *1* | *0.06* | 0.81 |
| | *sex* | *1* | *0.44* | *0.50* |
| | *Ancestral temperature* | *1* | *163* | *< 0.001 **** |
| | *sex x Lineage* | *1* | *1.05* | *0.30* |
| | *sex x ancestral temperature* | *1* | *0.95* | *0.33* |
| | *lineage x ancestral temperature* | *1* | *0.34* | *0.56* |
| | *sex x ancestral temperature x lineage* | *1* | *0.47* | *0.49* |

**Tab. S3:** *mean (± SE) O2 fluxes in picomol.s$^{-1}$ .mg$^{-1}$ according to the ancestral temperature (COLD C or WARM W) and assay temperature (25°C, 38.9°C, 43.8°C) for OCRs and ratios. Nechant corresponds to the number of tissue samples for each condition.*

| | Tissue | condition | Nechant | Mean | ± SE |
|---|---|---|---|---|---|
| LEAK CI | brain | C-25 | 21 | 9.22 | 1.21 |
| | | C-38.9 | 20 | 12.20 | 2.31 |
| | | C-43.8 | 18 | 10.75 | 1.83 |
| | | W-25 | 23 | 7.11 | 0.61 |
| | | W-38.9 | 20 | 15.59 | 1.17 |
| | | W-43.8 | 21 | 14.46 | 2.69 |
| OXPHOS CI | brain | C-25 | 21 | 12.78 | 1.55 |
| | | C-38.9 | 20 | 9.36 | 2.54 |
| | | C-43.8 | 18 | 5.56 | 1.49 |
| | | W-25 | 23 | 11.41 | 1.33 |
| | | W-38.9 | 20 | 15.95 | 1.72 |
| | | W-43.8 | 21 | 10,12 | 2.03 |
| OXPHOS CI+II | brain | C-25 | 21 | 15.85 | 2.23 |
| | | C-38.9 | 20 | 13.61 | 3.44 |
| | | C-43.8 | 18 | 9.99 | 2.38 |
| | | W-25 | 23 | 14.56 | 1.87 |
| | | W-38.9 | 20 | 19.12 | 2.98 |
| | | W-43.8 | 21 | 11.20 | 2.14 |
| ETS CI + II | brain | C-25 | 21 | 24.09 | 3.41 |
| | | C-38.9 | 20 | 14.49 | 3.85 |
| | | C-43.8 | 18 | 9.31 | 2.25 |
| | | W-25 | 23 | 20.97 | 2.99 |
| | | W-38.9 | 20 | 20.12 | 3.81 |
| | | W-43.8 | 21 | 10.06 | 2.03 |
| COX | brain | C-25 | 21 | 55.42 | 7.3 |
| | | C-38.9 | 20 | 112.95 | 14.88 |
| | | C-43.8 | 18 | 130.20 | 18.49 |
| | | W-25 | 23 | 44.87 | 5.56 |
| | | W-38.9 | 20 | 76.69 | 10.40 |
| | | W-43.8 | 21 | 77.00 | 14.29 |
| RCR.CI | brain | C-25 | 21 | 1.44 | 0.12 |
| | | C-38.9 | 20 | 0,77 | 0.08 |
| | | C-43.8 | 18 | 0.59 | 0.10 |
| | | W-25 | 23 | 1.65 | 0.19 |
| | | W-38.9 | 20 | 1.02 | 0.06 |
| | | W-43.8 | 21 | 0.68 | 0.07 |
| SCR | brain | C-25 | 21 | 0.44 | 0.06 |
| | | C-38.9 | 20 | 0.28 | 0.34 |
| | | C-43.8 | 18 | 1.92 | 0.74 |
| | | W-25 | 23 | 0.34 | 0.03 |
| | | W-38.9 | 20 | 0.53 | 0.13 |
| | | W-43.8 | 21 | 1.42 | 0.50 |

**Tab. S4:** *Analysis of Deviance Table (Type II Wald chisquare tests)* for LEAK CI, OXPHOS CI, OXPHOS CI+II, ETS CI+II and COX activity. When the interaction between predictors was significant we spliced the dataset by PREDICTORS variables.

| RESPONSE VARIABLE | PREDICTORS (Fixed effets) | DF | CHISQ | P |
|---|---|---|---|---|
| **LEAK-CI** | Lineage | *1* | *0.28* | 0.59 |
| | treatment | *1* | *0.005* | *0.94* |
| | Sex | *1* | *1.04* | 0.31 |
| | *Assay* | *2* | *16.5* | *< 0.001 **** |
| | *Ancestral* | *1* | *1.05* | *0.31* |
| | *Ancestral x Assay* | *2* | *4.9* | *0.09* |
| **OXPHOS-CI** | lineage | *1* | *0.12* | *0.74* |
| | treatment | *1* | *0.09* | *0.76* |
| | sex | *1* | *0.9* | *0.34* |
| | *Assay* | *2* | 14.1 | < 0.001*** |
| | *Ancestral* | *1* | *5.28* | *0.03** |
| | *Ancestral x Assay* | 2 | 9.94 | 0.006 ** |
| **OXPHOS-CI+II** | lineage | *1* | *1.30* | *0.25* |
| | treatment | *1* | *0.10* | *0.75* |
| | sex | *1* | *1.38* | *0.24* |
| | *Assay* | *2* | *17.2* | < 0.001 *** |
| | *Ancestral* | *1* | *1.08* | *0.30* |
| | *Ancestral x Assay* | *2* | *6.14* | 0.05* |
| **ETS-CI+II** | lineage | *1* | *0.52* | *0.22* |
| | treatment | *1* | *0.14* | *0.7* |
| | sex | *1* | *0.75* | *0.38* |
| | *Assay* | *2* | *46.33* | < 0.001 *** |
| | *Ancestral* | *1* | *0.36* | *0.55* |
| | *Ancestral x Assay* | *2* | *5.93* | *0.051* |
| **COX-activity** | lineage | *1* | *0.51* | *0.48* |
| | treatment | *1* | *1.03* | *0.31* |
| | sex | *1* | *2.04* | *0.15* |
| | *Assay* | *2* | *23.4* | < 0.001 *** |
| | *Ancestral* | *1* | *11.5* | < 0.001 *** |
| | *Ancestral x Assay* | *2* | *3.48* | *0.18* |

**Tab. S5:** *Analysis of Deviance Table (Type II Wald chisquare tests)* for RCR.CI and SCR variables. When the interaction between predictors was significant we spliced the dataset by PREDICTORS variables.

| RESPONSE VARIABLE | PREDICTORS (Fixed effets) | DF | CHISQ | P |
|---|---|---|---|---|
| **RCR.CI** | Lineage | *1* | *4.73* | *0.03** |
| | treatment | *1* | *0.10* | *0.75* |
| | Sex | *1* | *0.04* | 0.84 |
| | *Assay* | *1* | *94.2* | *< 0.001 **** |
| | *Ancestral* | *1* | *1.76* | *0.18* |
| | *Ancestral x Assay* | *2* | *0.95* | *0.62* |
| **SCR** | lineage | *1* | *0.38* | *0.54* |
| | treatment | *1* | *0.04* | *0.83* |
| | sex | *1* | *1.76* | *0.18* |
| | *Assay* | *2* | 15.5 | < 0.001*** |
| | *Ancestral* | *1* | *0.11* | *0.73* |
| | *Ancestral x Assay* | 2 | 1.13 | 0.57 |

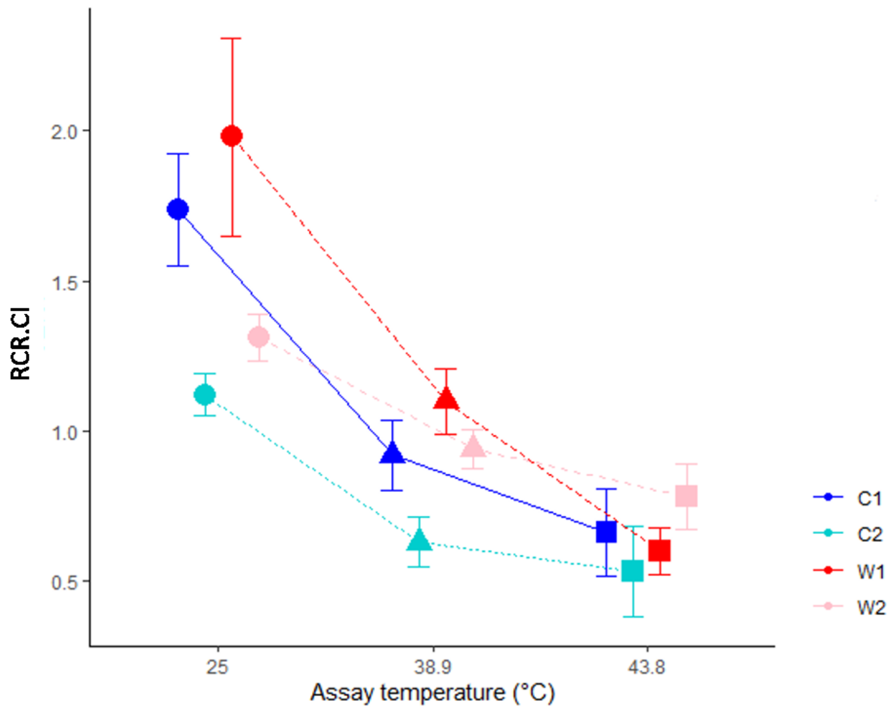


***Fig. S5***: *Mean (± SE) of RCR.CI ratio according to the ancestral temperature with the two lineages C1 & C2 for COLD and W1 & W2 for WARM fish (COLD 20°C fish C1 in blue; C2 in cyan; WARM 30°C fish W1 in red; W2 in pink) and assay temperature (25°C, dots; 38.9°C, triangles; 43.8°C, square).*

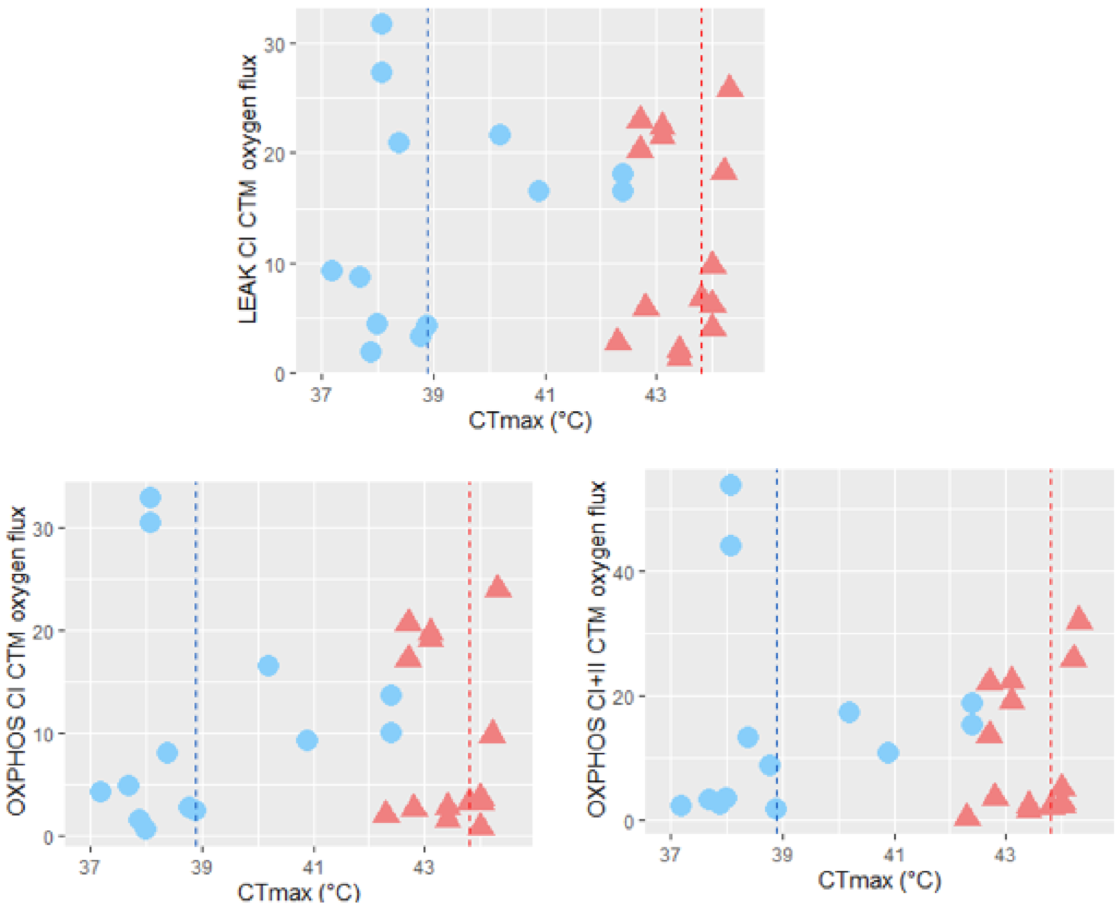


***Fig. S6:*** *OCR in picomol.s*$^{-1}$ *.mg*$^{-1}$ *at* $\overline{CTmax}$ *assay temperature for LEAK-CI, OXPHOS-CI, & OXPHOS-CI+II as a function of* $CT_{max}$ *whole-organism values for each thermal origin (COLD in blue circles or WARM in red triangles). Red and blue lines are* $\overline{CTmax}$ *assay temperatures for WARM and COLD fish, respectively.*

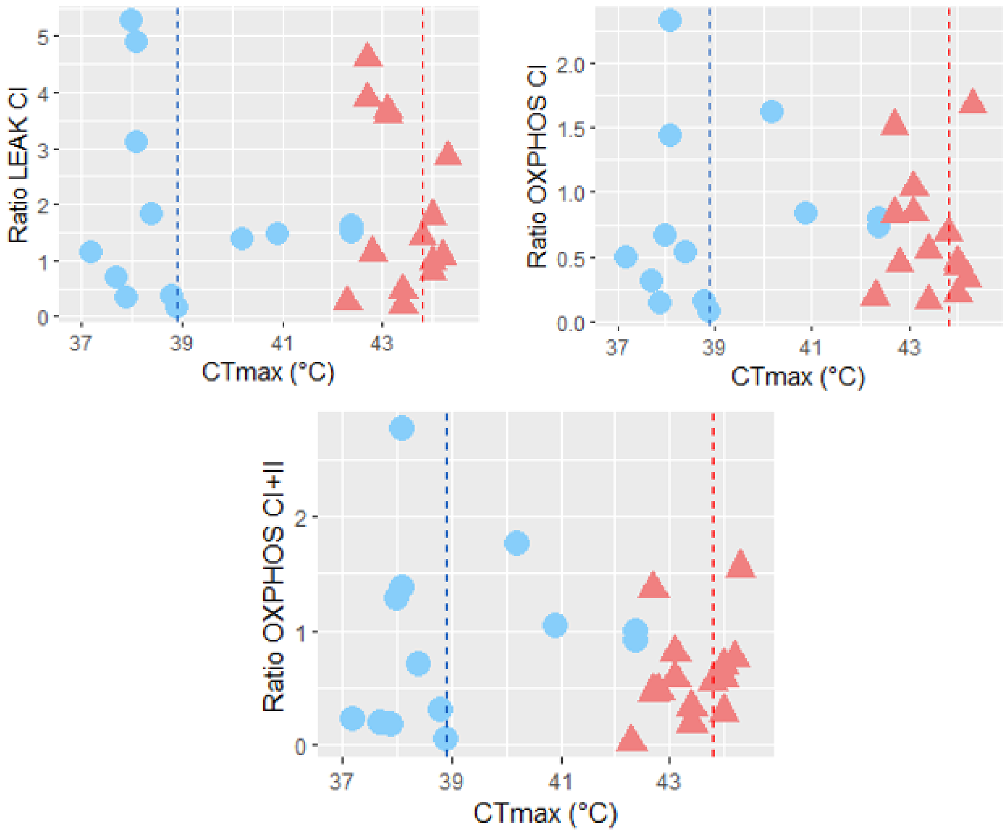


**Fig. S7:** *Ratio (OCR at $\overline{CTmax}$ / OCR at 25 °C) for LEAK-CI, OXPHOS-CI, & OXPHOS-CI+II as a function of* $CT_{max}$ *whole-organism values for each thermal origin (COLD in blue circles or WARM in red triangles). Red and blue lines are $\overline{CTmax}$ assay temperatures for WARM and COLD fish, respectively.*
